# A linear regression model predicts human brain ageing and reveals differential neuronal biological ageing relevant for Parkinson’s disease susceptibility

**DOI:** 10.64898/2025.12.01.691179

**Authors:** Shangli Cheng, Julio César Aguila Benitez, Melanie Leboeuf, Menghan Wang, Irene Mei, Silvia Gomez Alcalde, Qiaolin Deng, Rosario Sanchez Pernaute, Eva Hedlund

## Abstract

Biological brain ageing is a major risk factor for neurodegenerative diseases, which are characterized by selective degeneration of particular neuron types. We analyzed the impact of ageing on the transcriptome of neurons in the ventral tegmental area (VTA), substantia nigra pars compacta (SNc) and locus coeruleus (LC), that show differential vulnerabilities to Parkinson’s disease. Neurons were isolated from human *post mortem* brain tissues originating from 48 individuals ranging from 17 to 102 years of age and subjected to Smart-seq2 RNA sequencing. We identified 2,764 genes that were correlated with chronological ageing. This gene expression data was used to develop a feature selection-based Time Traversal algorithm, utilizing functionally grouped gene sets, GO terms, with high predictive accuracy of biological brain ageing. We identified 59 GO terms that can predict biological age using a linear regression model, where leave-one-out cross validation demonstrated a strong correlation between chronological age and predicted biological age (Pearson correlation coefficient = 0.946; adjusted R² = 0.771). The algorithm was validated on five independent datasets with high predictive performance, demonstrating shared ageing features across the human brain. Nonetheless, our analysis also highlights brain region and neuron type specificity in particular ageing features. Resilient neurons showed a weaker association with age-related transcriptional changes, indicating that they age slower than their vulnerable counterparts, thus revealing targets that may be used to slow down ageing and prevent disease development.

## Introduction

Chronological age is measured by the number of years since birth, whereas biological age is a measure of physiological health and a reflection of cellular damage accumulated over time. Individuals may age at distinct paces, and this can also change drastically as a result of diseases as well as interventions to counteract these. For instance, chronic diseases or specific medical treatments can cause biological ageing to accelerate beyond chronological age. Biological ageing can also be transiently induced by certain physiological and medical conditions, such as traumatic surgery, pregnancy, and in models, by heterochronic parabiosis (Poganik et al., 2023). Notably, parabiosis has been shown to temporarily reverse the biological age of the older individual, affecting factors such as blood vessel formation and adult neurogenesis (Boets et al., 2013; Sinha et al., 2014). Biological ageing is well recognized as a reliable predictor of health outcomes and is more accurate predictor, for instance, in predicting hospital mortality in critically ill individuals compared to chronological age (Ho et al., 2023).

For a long time, it was believed that longevity runs in certain families driven by genetic factors. However, it was recently demonstrated that the heritability of longevity is consistently low, less than 15%. Additionally, heritability estimates derived from genetic relatives also extend to nongenetic (in-law) relatives, suggesting that environmental factors, including life-style choices, play a major role in biological ageing, possibly alongside assortative mating (Graham Ruby et al., 2018). Biological age can be estimated through physical activity, performance metrics, medical records and imaging modalities (Manca et al., 2023).

Ageing is the primary risk factor for neurodegenerative diseases including Parkinson’s disease (PD), Alzheimer’s disease (AD) and amyotrophic lateral sclerosis (ALS). Understanding how ageing affects the brain may provide insight into why and how neurodegenerative diseases develop and why certain neurons are particularly vulnerable to the ageing process and degeneration. The difference between a person’s chronological age and their predicted brain age, the so-called brain age gap (BAG), determined by machine learning analyses of MRI scans, can be used to predict cognitive decline and dementia conversion, mostly related to AD. Furthermore, brain age estimated from plasma proteomics has recently been found to be one of the determinants of longevity (Oh et al., 2025). Studying how ageing impacts particularly vulnerable neuron types may help in designing therapies to target accelerated ageing of specific neurons, and thus render these more resilient to diseases such as PD. However, to conduct such analyses we need to ascertain that biological ageing can be reliably predicted within discrete neuronal populations. So how can we measure biological ageing in specific brain regions or neuronal subtypes and distinguish it from chronological age? Several studies have focused on evaluating and selecting markers to predict biological age and have employed different approaches, such as the regression based Klemera-Doubal method, principal component analysis (PCA), neural networks and multiple linear regression (MLR) (Husted et al., 2022)(Z. Li et al., 2023), machine learning (ML) and mathematical modelling (Xiong et al., 2023). In the MethBank database, a comprehensive DNA methylation repository for ageing research, ageing was categorized into twelve groups using age-specific methylated regions/cytosines (R. Li et al., 2018). ML models include both binary classifications, young versus old, as well as continuous prediction of ageing. In the mouse brain, a linear model successfully categorized samples as young or aged (Yu et al., 2023).

In this study, we used age-related Gene Ontology (GO) terms derived from RNA sequencing data as predicted features rather than the age-related genes themselves. In this way we could retain ageing related gene differences, while reducing the spread of individual samples with resulting increased prediction accuracy. We developed a linear regression (LR) model, using 59 of these GO terms, to predict biological brain age in three neuron types related to vulnerability in PD. We achieved a ‘Pearson’ correlation coefficient of 0.946 and an adjusted R squared of 0.771, using leave-one-out cross-validation (LOOCV). Predictions were highly correlated with chronological age across five independent brain datasets. However, our analysis revealed that while some ageing features are shared across neuron types, others are cell-type specific, with the strongest correlations observed in the cell type from which the data were derived. We also calculated the trajectories of 2,764 genes from the 59 Gene Ontology (GO) terms to identify the regulatory patterns, which were validated in these datasets. Importantly, the three neuron types investigated, displayed differences in biological ageing rates which relate back to their differential vulnerabilities and demonstrate that vulnerable neurons appear older than the chronological age of their host as well as of their more resilient neighbors. In conclusion, these findings demonstrated that analysis of age-related transcriptomic changes can reveal disease relevant pathways and targets.

## Results

### Identification of neuronal ageing-related GO terms through RNA sequencing of *post mortem* tissues spanning nine decades

To gain insight into how different neurons respond to the ageing process over a lifetime, we used laser capture microdissection (LCM) to isolate neurons from human *post mortem* samples of individuals age 16 to 102 years, followed by Smart-seq2 RNA sequencing (LCM-seq) (Nichterwitz et al., 2016, 2018). We hypothesized that neurons that show differential vulnerability to ageing-related-diseases such as PD may respond uniquely to the ageing process and thus show distinct biological ageing trajectories. Consequently, we conducted LCM-seq on; *i*) dopamine neurons from the ventral tegmental area (VTA), which are relatively resilient to PD (and potentially also to ageing), *ii*) dopamine neurons from the substantia nigra pars compacta (SNc), and *iii*) noradrenergic neurons from the locus coeruleus (LC), both of which are highly vulnerable to PD and ageing-related processes. We collected a total of 111 neuron samples originating from 48 brains, with each sample containing 25-150 neurons (**Table S1**).

The expression of specific neuronal marker genes and the absence of astrocyte, microglia and oligodendrocyte markers confirmed the neuronal identity of the LCM-seq samples (**Figure 1A**). Dimensionality reduction using Uniform Manifold Approximation and Projection (UMAP), based on expression profiles of previously identified markers of SNc and VTA (Aguila et al., 2021) demonstrated that VTA, SNc and LC neurons clustered separately, with LC showing the most distinct separation from SNc and VTA, as expected (**Figure 1B**).

**Figure 1.**
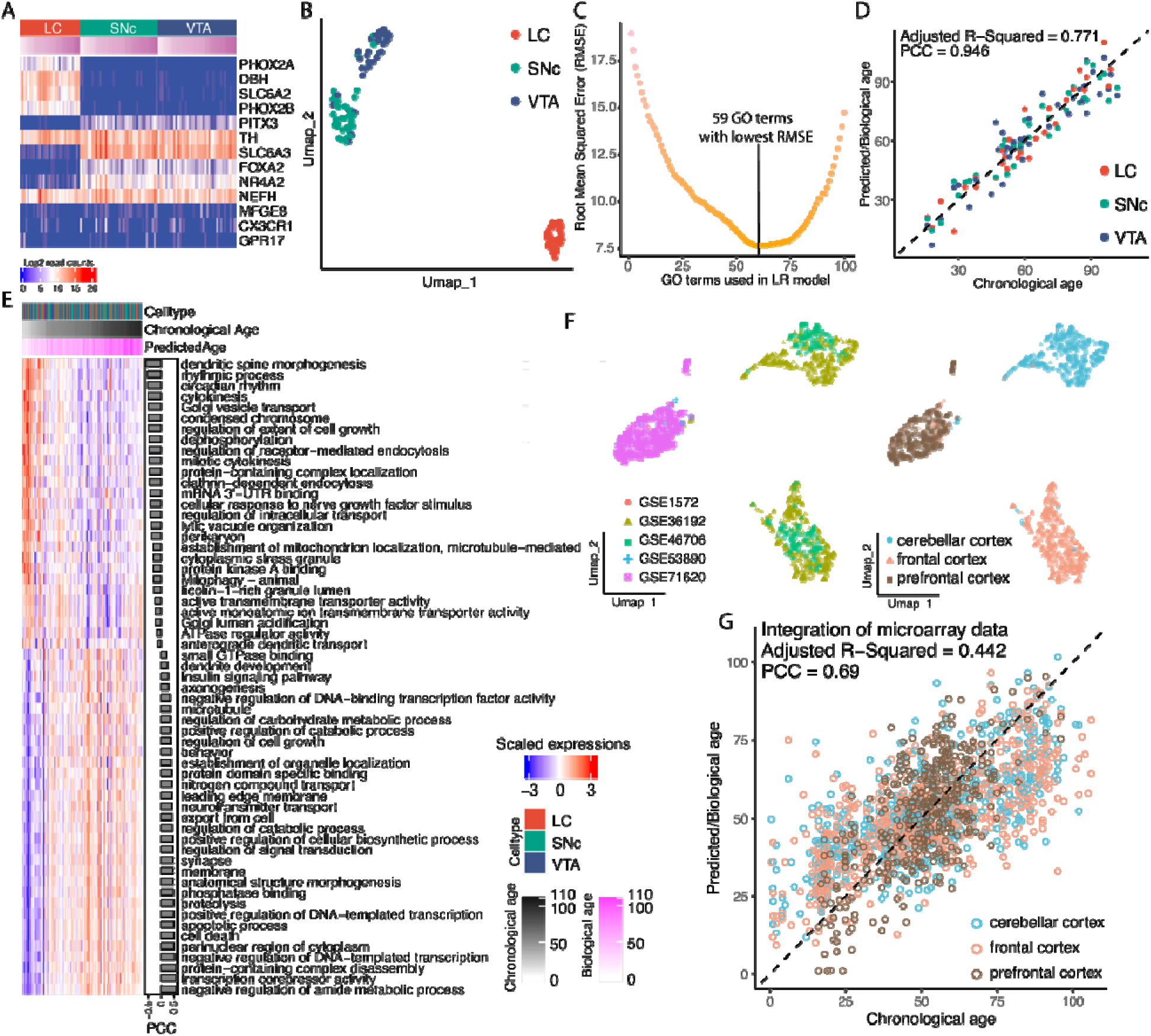
Feature selection and LR model in predicting biological age. **A**) Heatmap of marker gene expression in LC, SNc and VTA. **B**) Dimension reduction of UMAP with highly variable genes to separate LC, SNc and VTA. **C**) Feature-RMSE plot shows the increase of root mean squared error (RMSE) by the number of the feature GO terms in LR model. **D**) The predicted biological age is highly correlated with the chronological age. The adjusted R-squared is corrected by the number of the features. The Pearson correlation coefficient (PCC) between the chronological and predicted age was 0.946, and the adjusted R-squared was 0.771. **E**) The mean expressions of the genes from the 59 feature GO terms. The samples were ordered by the biological age from left to right. The side-by-side barplot is the ‘Pearson’ correlation coefficients between the mean expressions and the chronological age. The heat maps showing LC, SNc and VTA separately are provided in Figure S8. **F**) The integration of the five microarray data sets from human brain tissue and dimension reduction of UMAP colored by five data sources (left) and three sample types (right). **G**) Scatter plot shows the predicted biological age and chronological age in the integrated five data sets from the LR model using the 59 feature GO terms.

To identify ageing-associated GO terms, we first calculated the Euclidean distances between any two genes throughout the 111 samples by using their normalized expression the DESeq2 R package (version 1.44.0). The genes were then hierarchically clustered by using the “Ward.D2” agglomeration method (Murtagh & Legendre, 2014). Next, we computed the ‘Pearson’ correlation coefficients (PCC) of the gene clusters between chronological age and the mean expressions of the genes within each cluster across the LC, SNc and VTA samples. Clusters with PCC > 0.3 or < -0.3, indicating a moderate to strong (direct or inverse) correlation with ageing (Shaun Turney, 2024), contained 3,537 genes in 904 GO terms, enriched by GO term enrichment analysis with the g:Profiler (R package ‘gprofiler’ version 0.2.3) (**Table S2**). These ageing-related GO terms exhibited a statistically significant correlation with human brain ageing (**Figure S1A-C**). We calculated the sample variance, which measures the dispersion or spread of data and the results showed that the population variances characterized from GO terms were significantly less than those characterized from genes (**Figure S1D-E**). This indicates that using the GO terms we could capture the ageing differences while decreasing sample variability, which may improve the accuracy of the prediction model. Therefore, we selected GO terms as candidate features for further analyses.

### Feature selection and brain biological age prediction

To predict the biological age of the brain regions with a LR model, we employed the ageing-associated GO terms as features. The mean expression of the genes within each GO term was used as input to train and test the LR model, which was created using the R package ‘*caret*’ (Kuhn, 2008). To select an optimal set of GO term features, we developed a novel feature selection method, the *Time Traversal Algorithm*, which, together with a feature-RMSE (root mean squared error) plot, allowed us to evaluate and decide both the importance and ideal number of features (**Figure S2**, details in **Method**). The order in which features were selected reflects their contribution to modeling biological age. Given that our dataset contained 111 samples, we limited the feature-RMSE analysis to fewer than 100 features. To avoid overfitting and ensure unbiased performance estimation, we applied LOOCV to predict biological age. Model performance was evaluated using PCC between predicted biological and chronological age, as well as adjusted R-squared. Ultimately, a set of 59 GO terms yielded the lowest RMSE (**Figure 1C, Table S3**). When pooling the LC, SNc and VTA sample, the PCC between chronological and predicted biological age was 0.946, and the adjusted R-squared, normalized by the number of features, was 0.771. The unadjusted R-squared was 0.894 (**Figure 1D**).

Among the 59 feature GO terms, eight GO terms related to neuronal development and activity, including ‘dendritic spine morphogenesis’, ‘cellular response to nerve growth factor stimulus’, ‘perikaryon’, ‘anterograde dendritic transport’ were down-regulated with ageing. Moreover, two clearly detrimental GO terms, ‘apoptotic process’ and ‘cell death’, were up-regulated with ageing. In addition, two GO terms associated with Golgi regulation (Golgi vesicle transport and Golgi lumen acidification) were enriched. Interestingly, ‘cytoplasmic stress granules’ were found to be downregulated in neurons of older individuals, a change that has been linked to neurodegeneration, as stress granules decrease stress-related damage (Mahboubi & Stochaj, 2017). On the other hand, ‘dendrite development’, ‘neurotransmitter transport’, ‘synapse and axonogenesis’, were up-regulated with ageing, suggesting the existence of compensatory events occurring within neurons (**Figure 1E**). In conclusion, the GO terms associated with neuronal ageing reflect both the decline of key processes essential for neuronal integrity and function, and the activation of degenerative as well as compensatory protective mechanisms.

### Validation of the 59 feature GO terms using independent brain datasets

To validate the 59 feature GO terms selected for our LR model using the combined SNc, VTA and LC LCM-seq data, we collected five published brain microarray datasets with a total of 1,659 samples from three brain regions: cerebellar cortex (586 samples), frontal cortex (653 samples), and prefrontal cortex (420 samples) (**Table S4 Microarray datasets**). We first ensured the validity and consistency of the sample annotations by integrating all five datasets using ‘ComBat’ for batch correction (R package sva, version 3.52.0). Next, we performed dimensionality reduction of the PCA based on the 1,000 most highly variable genes and UMAP with the first ten PCs, which demonstrated that samples clustered according to brain region rather than dataset sources or potential batches (**Figure 1F**).

Our 59 GO terms were enriched from 2,764 genes, while there were 1,669 genes found in the five batch-corrected microarray datasets. To validate both the efficiency and robustness of the GO terms in prediction, we calculated the mean expressions of the genes within each GO term based on the identified genes. Then, we re-created the LR model with the same parameters described previously. By pooling the prefrontal cortex, frontal cortex and cerebellar cortex, biological age predictions were generated through LOOCV. The prediction showed a high PCC between chronological and predicted biological age (PCC = 0.69; adjusted R-squared = 0.442) (**Figure 1G**), confirming the generalizability and robustness of our selected GO-term features across multiple brain regions and datasets. The decrease in prediction accuracy could be related to the absence of some sample-type specific features in the independent datasets, which is in agreement with our hypothesis of cell-specific ageing. In conclusion, we have designed an algorithm based on RNA sequencing data that can select the most fitting features and predict ageing broadly, across neuron groups and datasets.

### The LR model demonstrates that neuronal ageing is in part cell-type specific

We next set out to investigate region as well as cell-type specific ageing patterns. As the number of samples should be higher than the number of features in LR models, we started our investigation using the microarray data sets from either prefrontal cortex, frontal cortex or cerebellar cortex, as these contained a large number of samples. With LOOCV, the predicted biological age of the samples was calculated within each brain region, separately. The results indicate that brain region-specific LR models achieved higher PCCs between predicted biological and chronological ages. Specifically, the PCCs for cerebellar cortex, frontal cortex and prefrontal cortex were 0.785, 0.734 and 0.801, respectively (**Figure S3A-C**). Meanwhile, the adjusted R-squared values improved to 0.498, 0.445 and 0.514. Therefore, a LR model trained on data from an individual brain region provided greater accuracy for that region than a model trained on data originating from several brain regions. To investigate whether the improved prediction accuracy was primarily due to the sample size or due to region-specific biological signatures, we performed a random subsampling analysis. We randomly selected subsets of 100 ∼ 400 samples from each region (cerebellar cortex, frontal cortex, or prefrontal cortex) to train the region-specific LR models and predict biological age with LOOCV. This subsampling was repeated 2,000 times for each region. The resulting mean PCC values for the three brain regions were 0.727 (cerebellar cortex), 0.667 (frontal cortex), and 0.757 (prefrontal cortex) (**Figure S3D**). The prefrontal cortex consistently showed the highest predictive performance, followed by the cerebellar cortex, and finally the frontal cortex. This indicates that the predictive performance of these LR models is primarily determined by the brain region origin of the transcriptome data, rather than the sample size.

Our LCM-seq starting sample sizes for LC (31 samples), SNc (40 samples) and VTA (40 samples) were below the original 59 GO-term features, making it statistically inappropriate to use all features in the LR models of the individual neuron types. Therefore, we performed a feature selection from the 59 GO terms for each neuron type (LC, SNc, VTA), recalculating feature-RMSE plots to determine the optimal subset of features. Altogether, we selected 16, 23 and 13 GO terms for LC, SNc and VTA, respectively (**Figure S4A**). When predicting biological age separately within each neuron type using LOOCV, the PCCs between the predicted biological and chronological age were 0.962 for the LC, 0.916 for the SNc and 0.849 for the VTA, with adjusted R-Squared values of 0.839, 0.609 and 0.582, respectively (**Figure 2A**). In conclusion, our analysis strongly indicates that different brain regions age distinctly and that even closely related neuron types, such as SNc and VTA dopamine neurons show differences in ageing trajectories, with VTA dopamine neurons appearing to age slower than disease vulnerable SNc and LC neurons.

**Figure 2.**
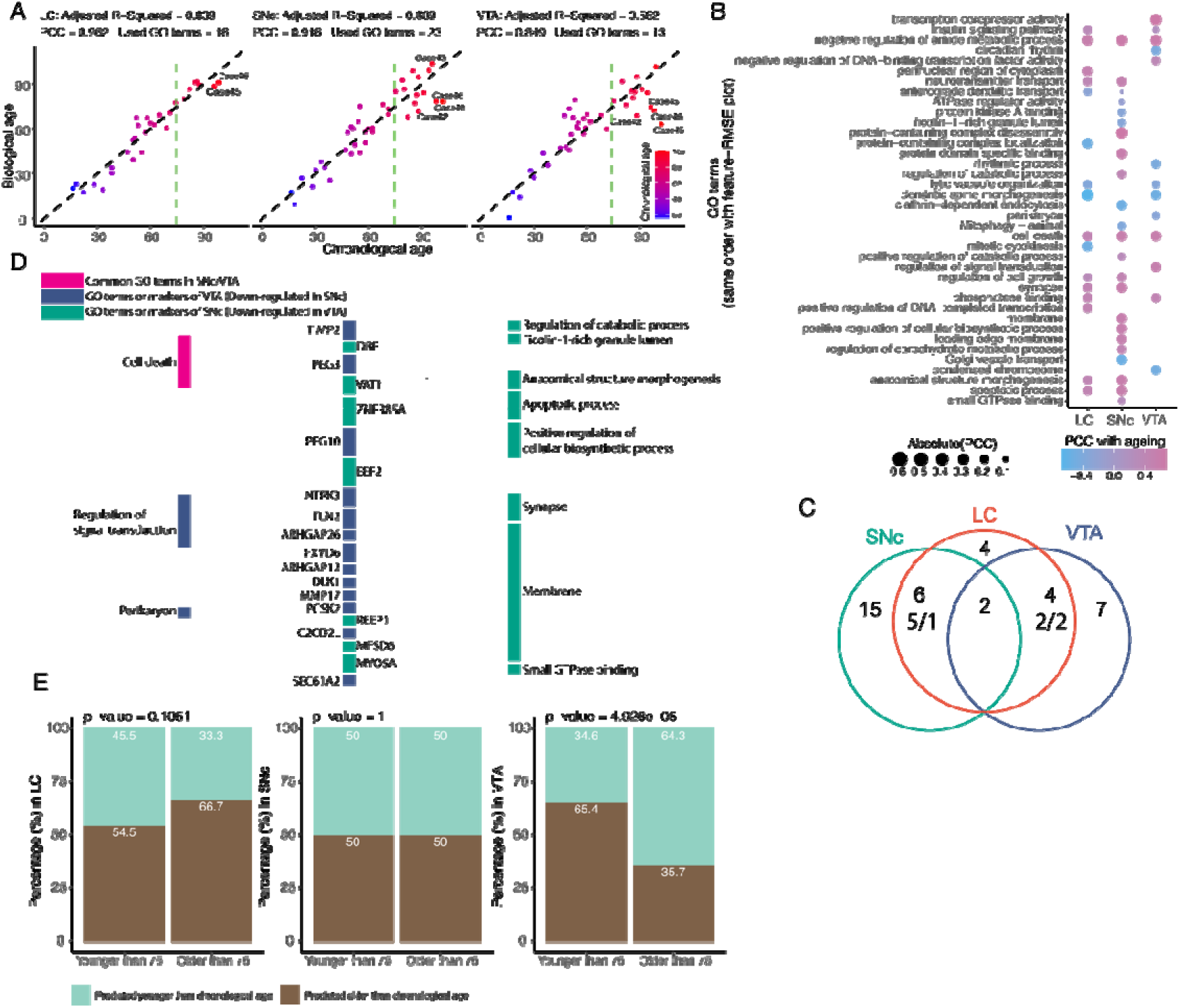
Sample type specific feature GO terms. **A**) LR models to predict biological age within LC, SNc or VTA. The number of feature GO terms used in the LR models of LC, SNc and VTA were 16, 23 and 13. The PCCs of LC, SNc and VTA between the chronological and predicted age were 0.962, 0.916 and 0.849, respectively. The adjusted R-squared values were 0.839, 0.609 and 0.582. The vertical dash lines separate the samples into two groups of younger and older than 75 (Figure S4B). **B**) The feature GO terms used in the three LR models of LC, SNc and VTA, where the size and color of the dots represent the PCC between the mean expressions of the GO terms with ageing. **C**) Venn diagram of the GO terms used in the 3 LR models of LC, SNc and VTA. There are 6 common GO terms between LC and SNc, in which 5 GO terms are up-regulated with ageing and 1 GO term is down-regulated with ageing. There are four common GO terms between LC and VTA, in which two GO terms are up-regulated with ageing and two GO terms are down-regulated with ageing. **D**) The Sankey plot shows the differentially expressed genes (DEG) between SNc and VTA and the GO terms including these DEGs. The mid-column is the DEGs and the right/left columns are the GO terms used in the LR model of SNc/VTA. **E**) The samples in Figure 2A were grouped into “younger” or “older” than 75 years of age. The bar plots show the normalized percentage of samples predicted “younger” or “older” compared to the chronological age. P-values are calculated from Chi-squared test.

### Feature GO terms reveal differences in biological ageing in LC, SNc and VTA neurons

Our LR models demonstrate that ageing involves both common molecular pathways and cell-type specific processes in LC, SNc and VTA. The feature GO terms used in each LR model differed between the three neuron types (**Figure 2B-C**). Interestingly, we found that SNc shared more features (GO terms) with LC than with VTA (**Figure 2C**). This suggests that despite their anatomical proximity and neurochemical as well as transcriptional similarity (**Figure 1B**), SNc and VTA neurons may age through distinct biological mechanisms, which could relate to their differential vulnerabilities. Only two GO terms were common to the three neuron groups: ‘negative regulation of amide metabolic process’ and ‘cell death’, both associated with the regulation of apoptosis.

Besides these shared terms, the LC and SNc models shared six GO terms, while LC and VTA shared four. Moreover, five out of the six common LC-SNc GO terms were upregulated with age, including ‘neurotransmitter transport’, ‘regulation of cell growth’, ‘synapse’, ‘anatomical structure morphogenesis’, and ‘apoptotic process’. These likely reflect both compensatory and degenerative responses during ageing. On the other hand, ‘anterograde dendritic transport’ was downregulated, pointing to impaired neuronal function. At the same time, among the four GO terms shared by LC and VTA, two were upregulated, including ‘insulin signaling pathway’ and ‘phosphatase binding’, while ‘lytic vacuole organization’ and ‘dendritic spine morphogenesis’ were down-regulated (**Figure 2B-C**). To find whether the specific GO terms in the SNc and VTA models reflected cell-type specific gene expression, we collected the GO terms including differentially expressed genes (DEGs) between SNc and VTA (Aguila et al., 2021). We found nine SNc GO terms and three VTA GO terms, with ‘cell death’ being the only shared term. In the Sankey plot, the VTA or SNc DEGs were positioned in the middle column, linking to either SNc-(right column) or VTA-specific (left column) GO terms (**Figure 2D**). All VTA markers (up-regulated in VTA and down-regulated in SNc) appeared in SNc GO terms, while only one SNc marker, ZNF385A, was found in one VTA GO term. Notably, PEG10 and PEG3, involved in regulating cell proliferation and apoptosis, were enriched in five and four GO terms, respectively. Some genes, such as PCSK2, a calcium dependent proteolytic enzyme which activates prohormones and neuropeptides, and C2CD2L (involved in insulin secretion and phosphatidylinositol transport) in the VTA, or VAT1 (vesicular transporter) in the SNc, are related to the differential aspects of neurotransmitter activity of these neurons. There were many genes in the GO terms enriched in VTA (relative to SNc) which relate to extracellular matrix (ECM) and cytoskeleton remodelling, including the youth associated protein-TIMP2 (Castellano et al., 2017), TLN2, MMP17 and the RHO-GTPases ARHGAP12 and ARHGAP26. Upregulation of these genes in the VTA may help counteracting aging-related changes, such as increased stiffness and disrupted synthesis, which can negatively impact synaptic function, trigger microglia activation and contribute to neurodegenerative diseases (Stern et al., 2022).

Moreover, compared to LR models of LC and SNc, the prediction of VTA had a lower PCC and adjusted R-Squared. Furthermore, when dividing samples into ‘adult’ and ‘old’ groups by 75-year old, we found that there were many more old samples predicted younger than their chronological age in the VTA, while the LC samples were more prone to be predicted older than their chronological age (**Figure 2E, S4B**). This result suggests that, in general, the VTA transcriptome is less affected by ageing than that of PD-vulnerable regions, and, particularly the LC, which is affected very early in AD and synucleinopathies (Matchett et al., 2021), and which appeared to display a marked ageing phenotype. These data support the notion that age-related transcriptomic changes may highlight targets driving neurodegeneration, and that cells and tissue that resist neurodegeneration also handle the ageing process better and may reveal genes and pathways relevant for rejuvenation.

### Gene trajectories of feature GO terms reflect the ageing process

As the 59 feature GO terms were enriched from 2,764 ageing genes, we next examined the regulatory and expression signatures of these genes. The 2,764 ageing genes were not associated with specific brain regions or functions; the initial two-dimensional PCA revealed no clusters among these genes, although ageing heterogeneity was observed in the PCA (**Figure S5A-B**). This suggests that the 2,764 genes are regulated in relation to brain ageing and do not exhibit sample type specificity. To further examine the regulatory and expression profiles of these genes, PCA was performed using the pooled set of 111 samples, projecting the 2,764 genes into lower dimensions (details in **Methods**). The 2-dimensional (2-D) coordinates of UMAP were calculated from the projected first 20 PCs. The principal curve based on the 2-D UMAP coordinates was subsequently calculated and taken as the order of the gene trajectories (**Figure 3A, S6A-B, Table S5**). We found that the gene trajectories were highly correlated with the ageing PCC (PCC of the 2,764 ageing genes) (**Figure 3B, S6C**). By ordering the 111 samples by age and averaging gene expression across adjacent groups of six samples, we identified genes with peak expression during ageing (see **Methods; Figure S6D**). Coloring the UMAP by age at maximum expression further revealed a correlation (PCC = 0.548) between expression signatures and gene trajectories **(Figure S6E**). These results indicate that gene trajectories capture both regulation and expression pattern information.

**Figure 3.**
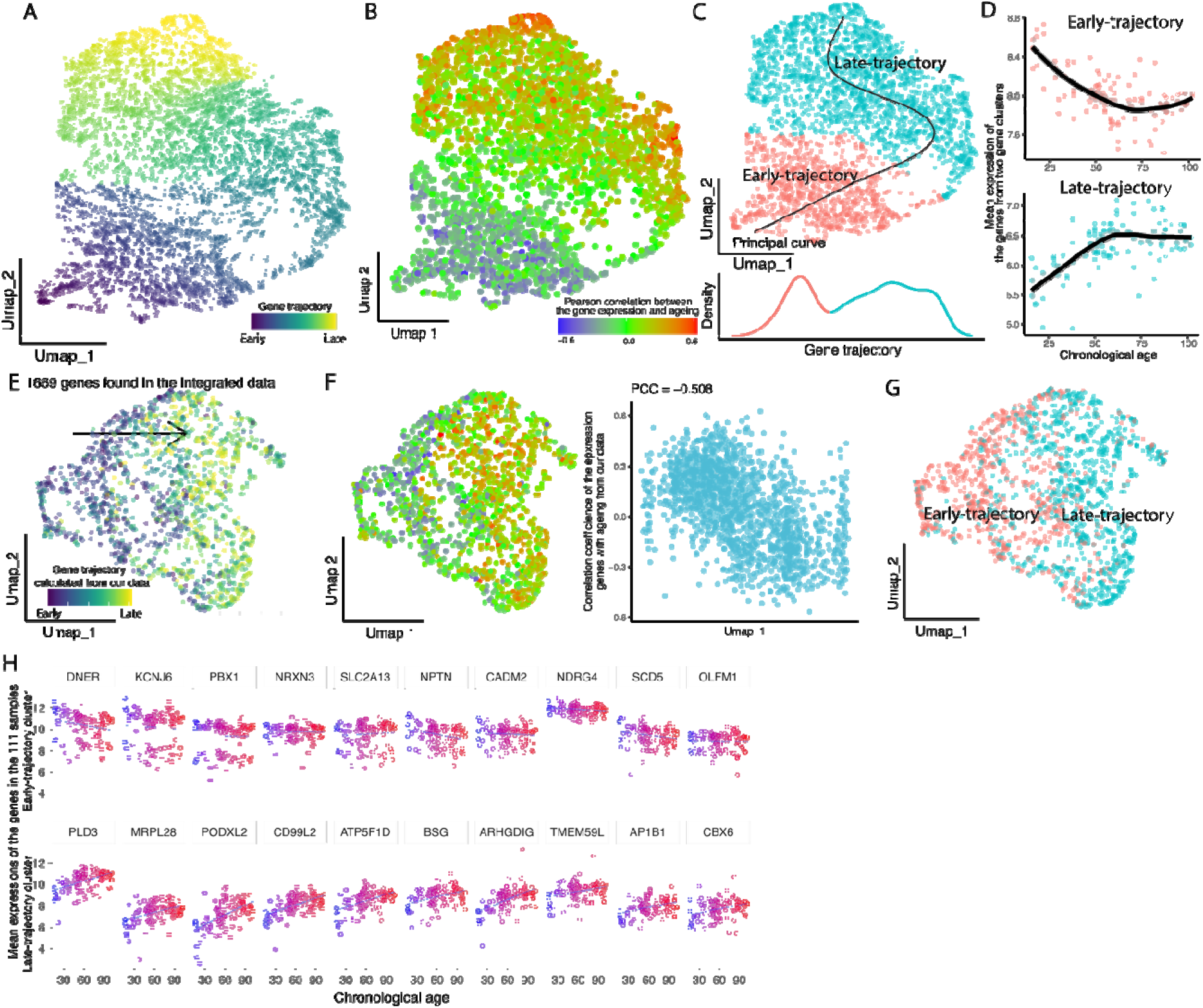
The trajectories of the genes from the 59 feature GO terms. **A-B**) Dimension reduction of UMAP show the trajectories of the genes from the 59 feature GO terms in A. The arrows follow the order of the genes calculated from principal curve. And the ‘Pearson’ correlation coefficients between the gene expressions and ageing were colored in B. **C**) The 2,764 genes from the 59 GO terms (up) were clustered into early or late trajectory clusters based on the density plot of the genes on the principal curve (down). **D**) By calculating the mean expressions of the genes from the two clusters of early/late-trajectory, the early trajectory genes show down-regulation, while the late trajectory genes show up-regulation with ageing. **E**) The UMAP of the 1,669 genes out of the 2,764 genes, which were detected in the five integrated microarray data sets. The color of the dots represents the gene trajectory values calculated as in Figure 3A. **F**) The ‘Pearson’ correlation coefficients between the gene expressions and ageing were shown in the five integrated data sets (left). And the genes show the pattern of up-/down-regulation with UMAP_1. **G**) The early and late trajectory genes colored as in Figure 3D. **H**) The expressions of the ten earliest and ten latest genes on the gene trajectory.

Furthermore, based on the density of the ageing genes along the principal curve, we grouped these into early and late trajectory clusters (Figure 3C). By calculating the mean expression of the genes in the two clusters, it becomes apparent that early trajectory genes tend to be down-regulated with ageing while highly expressed during youth, whereas late trajectory genes were up-regulated with ageing and showed lower expression in younger brains (**Figure 3C-D**). To test whether gene trajectories were dependent on the 59 GO terms, we averaged gene coordinates within each term to calculate the GO term trajectories. Most down-regulated GO terms clustered in the early trajectory region of the UMAP, while up-regulated terms appeared in the late trajectory area. This indicates that the GO terms reflect the trajectory patterns of their enriched genes and can be grouped into early or late clusters (**Figure S6F**).

To validate the gene trajectory of the ageing genes, we performed the PCA and UMAP analyses on the integrated five microarray datasets. Firstly, we found 1,669 common genes out of the 2,764 ageing genes, which were expressed in at least two samples. Secondly, the 2-D UMAP of the common genes, show similar trajectories to Figure 3A, with early- to late-expression, from left to right (**Figure 3E**). Moreover, we also colored the UMAP by the ageing PCC (same values as in Figure 3B) and the expression signatures (same values as in Figure S6E) (**Figure 3F and S6G**) and found that the ageing PCC showed a high correlation (R = 0.508) with UMAP_1 (**Figure 3F**). In addition, early and late trajectory clusters were identified within the 1,669 genes, with early trajectory cluster genes primarily found on the left, while the late trajectory cluster genes were mainly on the right (**Figure 3G**).These findings confirm the trajectories of the ageing genes across independent datasets.

### Identified neuronal ageing genes demonstrate improved accuracy in predicting brain ageing than general ageing genes

Previous studies have reported ageing genes in three ageing genomic databases, Human Ageing Genomic Resources (HAGR) (de Magalhães et al., 2005), Ageing Atlas (AA) (Liu et al., 2021) and AgeingReG (Piao et al., 2023). As the ageing gene databases include non-tissue-specific ageing genes while our ageing genes are brain-specific, there are only 16 ageing genes in common between these three data bases and our data set of LC, SNc and VTA (**Figure 4A**) of which seven, ATM, GSK3b, HDAC2, HMGB1, PTEN, SOD1 and XRCC5 were from the early trajectory cluster and nine genes, AKT1, CLU, FOXO3, GSK3A, LMNA, SIRT6, SIRT7, SP1 and VEGFA from the late trajectory cluster (**Figure 4B**).

**Figure 4.**
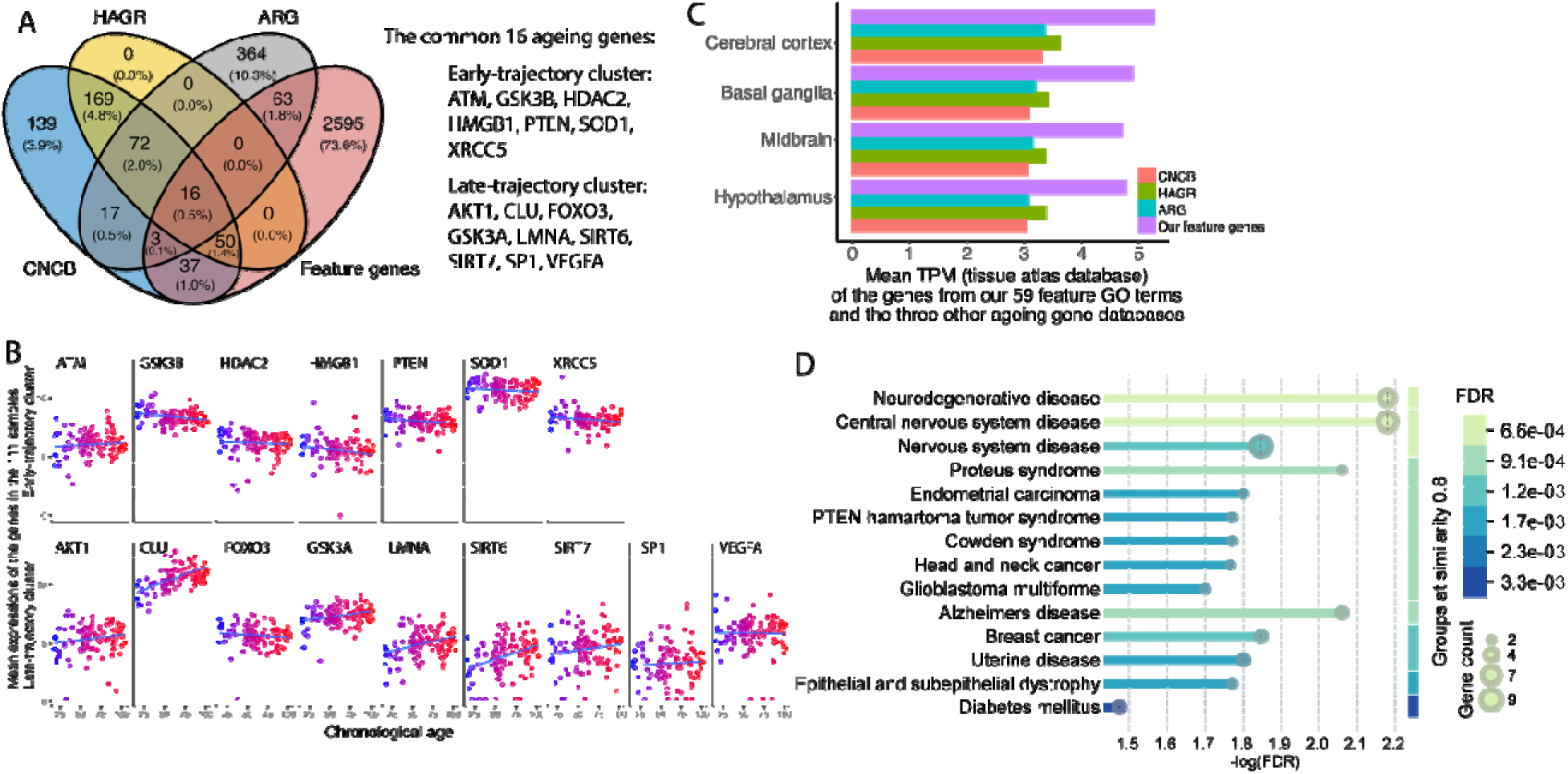
Comparisons between our ageing genes with other ageing gene databases. **A**) Venn diagram shows the comparisons between the 2,764 genes from the 59 feature GO terms and the other three ageing gene databases (left). **B**) The expressions of the 16 overlapping ageing genes in Figure 4A. Out of the 16 genes, seven are from early-trajectory cluster and nine are from late-trajectory cluster (right). **C**) Based on the Consensus Transcript Expression levels summarized per gene (CTExp), the bar plot shows the mean expression of the 2,764 genes and the ageing genes from the three databases in the four brain tissues, including cerebral cortex, basal ganglia, midbrain and hypothalamus. **D**) Disease-gene Associations (DISEASES) enrichment in STRING by using the 16 common ageing genes with the ageing databases.

Moreover, to compare the expression levels in brain tissues of the ageing genes from our study and the three databases, we used Consensus Transcript Expression levels summarized per gene (CTExp) in 50 tissues based on transcriptomics data from HPA and GTEx. We extracted the normalized TPM of the ageing genes in brain tissues of cerebral cortex, basal ganglia, midbrain and hypothalamus from the CTExp expression matrix. The mean TPM of the genes from the three databases and our study were calculated in each tissue (**Figure 4C**). The expression of our ageing genes was higher than those from the three ageing gene databases in the four brain regions, demonstrating that brain specific gene regulation is needed for accurate prediction of brain biological ageing features. To further comprehend the overlapping 16 ageing genes throughout the different studies, we employed Disease-gene Associations (DISEASES) enrichment in STRING and found that most of the overlapping 16 genes were enriched in ‘nervous system disease’, ‘central nervous system disease’ and ‘Neurodegeneration disease’ (**Figure 4D**). In conclusion, our analysis shows that ageing-related is a tissue and neuron-type specific process and that the ageing genes and GO terms identified in our study are reliable predictors of brain ageing in general. Our approach identified processes of neuron degeneration with age and demonstrated that neuron resilience and susceptibility to ageing-related disease is correlated with their predicted biological ages, distinct from the chronological age.

## Discussion

In this study we analyzed 111 human brain samples (LC, SNc, and VTA) to examine gene expression patterns and their correlation with ageing. By leveraging gene expression data and focusing on functionally grouped gene sets—namely, GO terms—we aimed to reduce individual sample variability and improve predictive accuracy of biological brain ageing. GO terms were enriched based on mean gene expression, which showed both positive and negative correlation with ageing. Using a novel Time Traversal feature selection algorithm, 59 GO term features were chosen for the LR model predicting biological age. LOOCV across pooled samples yielded a strong correlation (PCC = 0.946, adjusted R² = 0.771) with chronological age. Validation on five independent datasets confirmed high predictive performance (PCC = 0.687). Dimensionality reduction of 2,764 genes led to identification of gene trajectories, distinguishing early and late trajectory genes highly expressed in young or old individuals.

Previous studies predicted biological age using machine learning models based on DNA methylation, gene expression, or biomarkers, but often showed lower correlations. The biological age prediction by using CpG sites showed higher accuracy and PCC. However, the studies using the expressions of the biomarkers obtained lower accuracy and PCC compared with our results (**Table S6**). At the molecular level, multiple factors contribute to biological ageing, including telomere length, DNA methylation, DNA damage and mitochondrial function (Jylhävä et al., 2017)(Bell et al., 2012)(Heyn et al., 2012)(López-Otín et al., 2013). Molecular measures used to assess biological ageing include DNA methylation, as well as transcriptomics and proteomic analyses across different tissues and species (Xiong et al., 2023)(Qiu et al., 2023)(Aadahl & Jørgensen, 2003)(Lu et al., 2023). Of these, the DNA methylation clock has emerged as the premier tool for assessing biological age as methylation levels at specific CpG sites change predictably with chronological age (Gyenis et al., 2023; Horvath, 2013). Moreover, besides common ageing factors, biological age is influenced by genomic modifications, such as neddylation as well as transcriptional elongation and its regulation (Saurat et al., 2024)(Debès et al., 2023). So far, numerous genes and pathways related to (biological) ageing have been identified. Transcriptional stress caused by endogenous DNA damage is correlated with ageing and is associated with pathways such as p53 signaling and apoptosis (Gyenis et al., 2023). In human brain studies, genes such as *DLGAP1*, *SOX10*, *OLIG2* and *DLGAP1-AS1* have shown ageing related-changes, particularly related to dividing glial cells (Soreq et al., 2017) (Herring et al., 2022). In mouse studies, *Cfl*, *Rpl13a*, *Lars2*, *P4hb*, *Slc5a4b* and *Slc13a4* have been found to be differentially expressed with ageing, independent of tissue specificity (Zhang et al., 2021) (Yu et al., 2023). Our transcriptomic LR model, which leverages ageing-related GO terms, demonstrated superior predictive accuracy (PCC = 0.946, adjusted R² = 0.771) compared to most biomarker-based or gene-based models, and approached the performance of DNA methylation models.

Compared to LR models constructed using pooled multiple types of samples, LR models based on integrated microarray data with a single sample type demonstrated improvement in either PCC or adjusted R-squared for biological age prediction. This clearly demonstrates that there are brain region-specific ageing processes at play. Conversely, LR models using only SNc or VTA samples showed reduced PCC or adjusted R-squared values compared to those using LC samples. Analysis of RMSE values across different ages indicated that LC samples generally had lower RMSE values than SNc and VTA. For identical sample cases, such as Case 38 and Case 46, most RMSE values for SNc or VTA were higher than for LC, particularly among samples from older individuals. This suggests that LC samples, both young and old, exhibit greater robustness than SNc and VTA. Furthermore, it also indicates that VTA neurons, isolated from individuals 75 years and older appear somewhat younger than predicted from their chronological, suggesting that VTA neurons age slower than LC neurons.

Previous studies have identified numerous biomarkers. *CEP170B*, *PPP1CB*, *CUK1*, *ARHGAP1*, *SEB1*, *PUM1*, *NOVA1*, and *ATXN2* were reported to show lower variability in AD compared to controls (Saurat et al., 2024). In this analysis, *CEP170B*, *GUK1*, and *ARHGAP1* were positively correlated with ageing, while *PPP1CB*, *PUM1*, *NOVA1*, and *ATXN2* were negatively correlated with ageing. A recent preprint demonstrated that knockdown of *nova-1* in C.elegans resulted in increased lifespan and increased heat stress recovery (Adeyemi et al., 2025), indicating that a lower level of *NOVA1* identified in our study could be a protective response to ageing. However, *NOVA1* is also involved in regulating *hTERT* splicing and loss of *NOVA1* results in a shift to generation of non-catalytic telomerase isoforms which could result in telomere shortening (Ludlow et al., 2018). Lowered levels of *ATAXN2* may lead to mitochondrial dysfunction, which could lead to impairment in calcium homeostasis and enzyme activities (Meierhofer et al., 2016). Nonetheless, it could also lead to improvements in other context, for example a reduction in protein aggregation (e.g. in TDP-43 and its recruitment to stress granules (Becker et al., 2017)).

In mouse studies, *Cep170b* was up-regulated and Ppp1cb down-regulated. Additional body and brain ageing genes were identified within the feature GO terms. The body ageing gene *FOXO3* was down-regulated with ageing, whereas *LMNA* and *ATM* were up-regulated. Research has shown an association between *FOXO3* and human ageing, including a relationship between specific single-nucleotide polymorphisms (SNPs) of *FOXO3* and longevity (Park, 2024). OPA1, linked to neurological manifestations, has also been found to be down-regulated in older individuals (Rusecka et al., 2018).

Up-regulation of *LMNA*, *SIRT6* and *CLU* indicates activation of compensatory mechanisms in healthy neuronal ageing. SIRT6 is a NAD+-dependent histone deacetylase broadly associated with longevity. Clusterin acts as a molecular chaperone, preventing aggregation of proteins such as *APP*, *B2M*, and synuclein (Yuste-Checa et al., 2022). Both protective and risk variants of *CLU* exist for AD. Indeed, a mutation that reduces expression of the secreted clusterin isoform (sCLU), affecting *A*β and tau protein aggregation and clearance, is the third most significant genetic risk factor for late-onset AD (Harold et al., 2009; Lambert et al., 2009). On the other hand, lower *SOD1* would put a cell at risk as it could lead to increased oxidative damage and resulting inflammatory responses, and it leads to faster ageing phenotypes in model systems (Watanabe et al., 2014). We acknowledge that this study has limitations namely due to the sample size, with only 111 samples derived from three neuron types, and thus in the LR model we currently utilize a limited number of GO term features. Future plans involve developing an LR model with more GO term features, based on additional samples, in order to achieve higher accuracy.

## Conclusion

We have developed a systematic approach to distinguish biological age from chronological age at the transcriptomic level in specific neuronal populations and identify different age trajectories related to neurodegeneration. By leveraging gene expression data and focusing on functionally grouped gene sets, GO terms, we could reduce individual sample variability and improve predictive accuracy of biological brain ageing. This methodology combines feature selection through a novel algorithm with robust statistical modeling, enabling us to interrogate mechanisms of ageing across brain regions and neuron types. We were able to identify ageing mechanisms shared across neuronal regions and types as well as molecular signatures unique to particular neurons, which may underlie their differential vulnerabilities to disease. This approach provides a framework for uncovering critical pathways relevant to neurodegenerative disease risk and resilience and sets the stage for future studies targeting interventions at the level of specific neuronal subtypes. The features associated with GO terms demonstrated effectiveness for brain biological age prediction in published datasets and in our neuronal samples, revealing differences related to PD vulnerability. Importantly, age-related transcriptomic changes in vulnerable and resilient neurons may highlight targets for therapeutic approaches in neurodegeneration.

## Supporting information

Supplementary Information including Figures

## Acknowledgments

The authors acknowledge support from the National Genomics Infrastructure in Stockholm funded by Science for Life Laboratory, the Knut and Alice Wallenberg Foundation and the Swedish Research Council, and SNIC/Uppsala Multidisciplinary Center for Advanced Computational Science for assistance with massively parallel sequencing and access to the UPPMAX computational infrastructure. This work was funded by grants to EH from Parkinsonfonden (number 795/15; 910/16; 991/17; 1095/18; 1192/19; 1258/20; 1328/21; 1413/2022; 1558/24); The Swedish Research Council (2020-01049; 2024-02902), startup funding from the Department of Biochemistry and Biophysics at Stockholm University and by the InnoHK initiative of the Innovation and Technology Commission of the Hong Kong Special Administrative Region Government. And by funding to RSP from Instituto de Salud Carlos III (ISCIII) PI24/00707. Human post mortem tissues were kindly received from the Netherland’s Brain Bank (NBB) and from the NIH NeuroBioBank.

## Author contributions

Supervision and Conceptualization: EH. Investigation: SC, JA, ML, SGA, IM, MW, QD, RSP, EH. Data curation, Methodology and Software: SC, IM. Validation: SC, ML, JA. Visualization: SC, ML. Resources: EH, JA. Writing - Original Draft, Review & Editing: SC, JA, ML, SGA, IM, MW, QD, RSP, EH.

## Methods

### Ethics Statement

We have ethical approval to work with human post mortem tissues (Supplementary Tables S1) from the regional review board of Stockholm, Sweden (Dnr 2012/111-31/1; 2012/2091-32) and The Swedish Ethical Review Authority (Etikprövningsmyndigheten) (Dnr 2023-00280-01). Human fresh frozen tissues were retrieved from the Netherlands Brain Bank (NBB) and the NIH NeuroBioBank. The work with human tissues was conducted according to the Code of Ethics of the World Medical Association (Declaration of Helsinki).

### Sample isolation and RNA-Seq

Neurons were isolated from post mortem tissues from individuals that were never diagnosed with a neurodegenerative disease and thus considered controls in this aspect, ranging from age 16 up to 102 years-of-age.

### Dimension reduction of the 111 samples

The principal component analysis (PCA) was based on the substantia nigra pars compacta (SNC) and ventral tegmental area (VTA) biomarkers (Aguila et al., 2021). Then, the UMAP of the 111 samples was calculated from the first 10 PCs (**Figure 1B**).

### Highly correlated genes with ageing and gene ontology enrichment

To find the expression patterns of the genes with ageing, the 23,699 genes were clustered into 200 clusters. The mean expression of the genes within each cluster was calculated. Then after, the ‘Pearson’ correlation coefficient between the mean expression of each gene cluster and ageing vector was calculated. The genes from the cluster with Pearson correlation coefficient higher than 0.3 (positively correlated) or lower than -0.3 (negatively correlated) were kept and used as highly correlated genes with ageing. Then, the gene ontology (GO) enrichment found the GO terms with positively correlated and negatively correlated genes separately by using g:Profiler (R package gprofiler2 version 0.2.3). There are 573 GO terms and 13 KEGG pathways were enriched with the positively correlated genes, while 645 GO terms and 25 KEGG pathways were enriched with the negatively correlated genes. For there are 352 common GO terms between the positively correlated GO terms and negatively correlated GO terms, we combined the common GO terms. Finally, 904 GO terms were obtained (**Table S2**).

### Compare Enriched ageing GO terms with GO database

To compare the positively and negatively correlated GO terms with the GO database, we also collected all GO terms from the human GO database (R package ‘org.Hs.eg.db’ version 3.19.1) (Marc, 2019). We analyzed Pearson correlation coefficient between the mean expression of the genes from each GO term and chronological age. There are 15,924 Pearson correlation coefficients from the same number of GO terms (**Figure S1A**).

### Time Traversal Algorithm to order the features

To evaluate the significance of the features for the linear regression (LR) model in prediction, we developed a novel feature selection algorithm, Time Traversal Algorithm (**Figure S2**):

1. there are N P-terms taken as the features for the LR model.
2. to select 1 P-term from N P-terms.

2.1) to create an LR model using the 1 P-term.
2.2) to calculate predicted biological age by leave-one-out cross validation.
2.3) to calculate the root mean square error (RMSE) between predicted biological age and chronological age, where RMSE measures the average difference between values predicted by a model and the actual values.
3. repeat (2) until all P-terms were selected.

3.1) to find the lowest RMSE and take the P-term as the first P-term/feature.
4. there are N-1 P-terms left.
5. to select 1 P-term from N-1 P-terms.
6. to repeat (2) and (3), find the second P-term/feature.
7. to repeat (5) and (6) until selecting the M number of P-terms.

Here we took M less than 80% of samples to avoid overfitting of the LR model.

The RMSE is defined as:

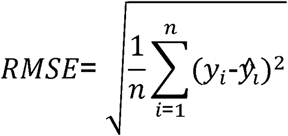

Where *y_i_* is chronological age, and *ŷ_i_* is predicted biological age.

### Evaluation method of leave-one-out cross-validation (LOOCV)

To evaluate the performance of an LR model on one dataset, we employed LOOCV, taking one sample as the validation test dataset and the remaining samples as the training dataset:

1. *input of N samples*.
2. *taking 1 out of N samples as the test sample and remaining N-1 samples as the training samples*.

*2.1) create LR model with N-1 training samples*.
*2.2) calculate predicted value of 1 test sample*.
3. *repeat (2), until N samples are all taken as test samples*.

### Feature-RMSE plot

To select and define the best features used in LR model based on the order of the GO terms calculated by Time Traversal Algorithm, we developed Feature-RMSE plot in which x-axis shows the numbers of the GO terms adding to the LR model and y-axis is the RMSE between the predicted age and chronological age. By adding more GO terms in order to LR model, RMSE will reach a local minimum. For RMSE can measure the average differences between predicted biological age and chronological age, simultaneously we got the minimum difference in prediction. Finally, we selected the GO terms as the final GO terms used in the LR model for biological age prediction.

### Linear regression model

The linear regression model was created and trained by using R package ‘caret’ and ‘train’ function with the formula of ‘Chronological age ∼ Features’ and the fitting method of ‘lm’.

### Evaluation of accuracy of linear regression model

Because the predicted values from linear regression are not measured by classifications, we cannot calculate accuracy to evaluate LR model. Alternatively, we employed Pearson correlation coefficient (PCC) and adjusted R-Square (R^2^) to evaluate the accuracy of LR model:

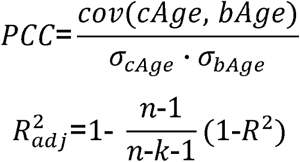

Where cAge is the chronological age; bAge is the predicted biological age; s is the standard deviation; *n* is the sample size and *k* is the number of predictor variables, here *k* is the number of GO terms used in LR model.

### Expression signature of the genes (Figure S6D)

To characterize the age when a gene reaches its highest expression in the 111 samples, we first ordered the samples by chronological age from 16 to 102 years of age. Then, the expression of a gene of each sample was averaged by the expressions of the nearest former and following six samples. Finally, we got the age when a gene achieve its highest expression (or maximum value in the expression array) and then ordered the genes (**Figure S6D**).

### Testing datasets from published studies

To validate the power of the 59 feature GO terms in predicting biological age for other human brain data, we collected five high-throughput microarray datasets of human brain tissue from published studies and one single-cell RNA-seq dataset. And we downloaded all datasets from Gene Expression Omnibus (GEO) (**Table S4**).

### Integration and dimensionality reduction of the five brain microarray datasets

To integrate the five microarray datasets, we corrected the expression matrices using the method of ComBat in R package ‘sva’ (version 3.52.0).

The PCA analysis was based on the 1,000 most variable genes. Finally, the UMAP was calculated from the first 10 PCs.

### Gene trajectories to characterize the expression patterns with ageing

Usually, we calculated PCA of the 111 samples (observations) using the expressions of 2,764 genes (variances). To recognize and characterize the gene expressions with ageing, we collected 2,764 genes (taken as observations) from the 59 feature GO terms. Next, we calculated the PCA of the genes based on 111 samples of LC, SNc, VTA (taken as variances). The first 20 PCs were projected into the 2-dimension of UMAP. Then, we calculated the principal curve from the coordinates of the first two dimensions of the genes in UMAP. Finally, we took the order of the genes as the gene trajectory.

### The ageing ‘Pearson’ correlation coefficient (PCC) of the genes and the gene expression signature

To calculate the ageing PCC of the genes, which represent the gene expression pattern with ageing, we first ordered the samples from young to old. Then for each gene, there would be a PCC between the expression and ageing vector.

To define the gene expression signature, firstly, we ordered the samples by chronological age from young to old. Secondly, the expression of a gene for each sample was averaged by the former and following six samples. Finally, the gene expression signature was defined as the age at the maximum expression.

### Statistics

In Figure 2E, we examine two categorical variables using ‘Chi-squared test’: 1) the percentage of the samples predicted younger and older than chronological age; 2) the groups of ‘younger than 75’ and ‘older than 75’.

