## Supplementary Information including Figures for "A linear regression model predicts human brain ageing and reveals differential neuronal biological ageing relevant for Parkinson’s disease susceptibility"

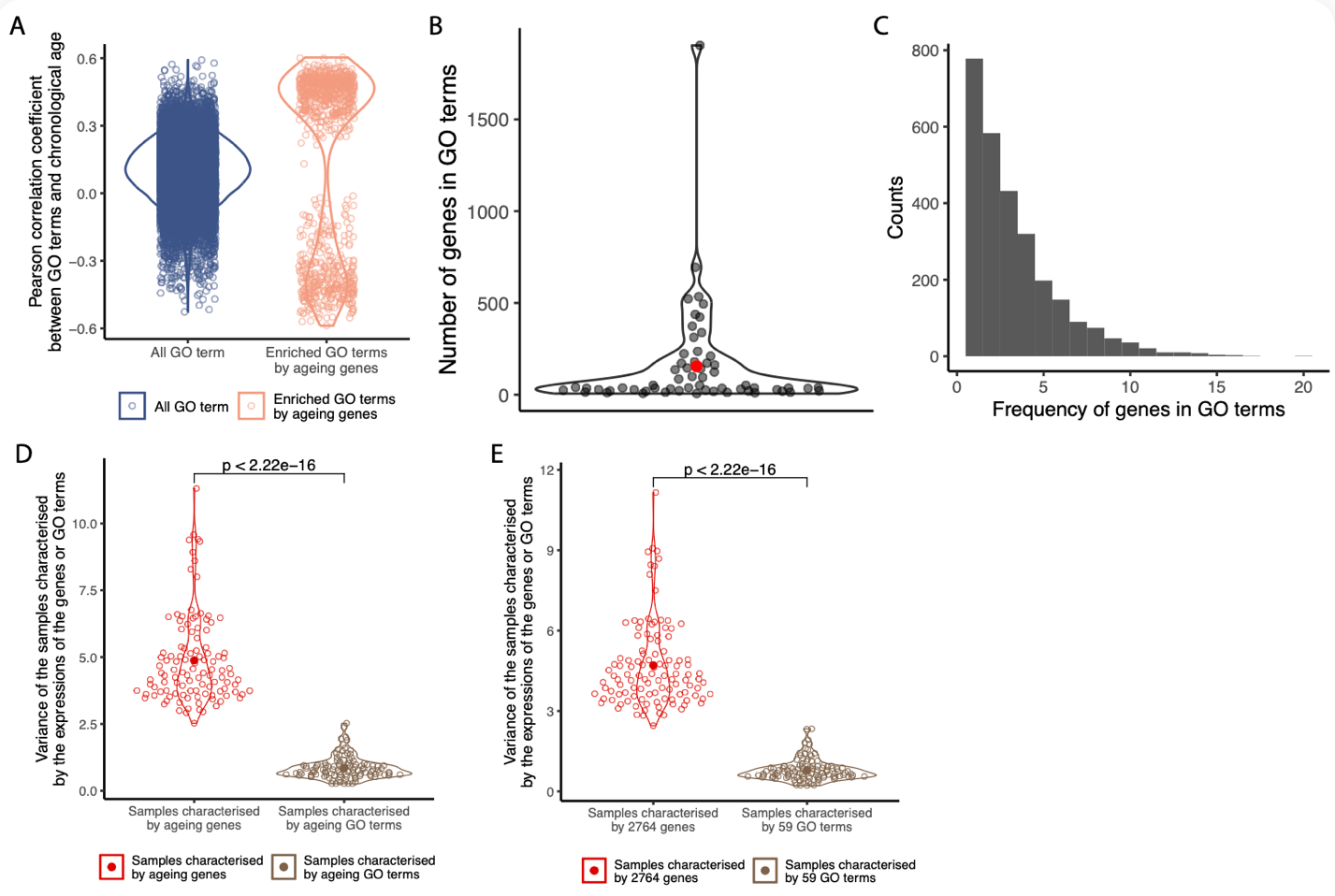

**Figure S1 The enriched GO terms**

A) Violin plot shows the Pearson correlation coefficients between the mean expressions of the genes of the GO terms and chronological age in all GO terms and the enriched GO terms from ageing genes. The P value of Wilcoxon test between the two conditions of the absolute PCC values is <0.001.

B) The numbers of the genes enriched in the 59 feature GO terms. The filled red dot shows the mean number.

C) One gene could be enriched in multiple GO terms. The histogram shows the frequency of the 2,764 genes enriched in the 59 feature GO terms.

D) The violin plot shows the variance of the samples calculated from the expressions of the ageing genes and GO terms.

E) The variance of the samples calculated from the 59 feature GO terms and the 2,764 genes out of the 59 GO terms.

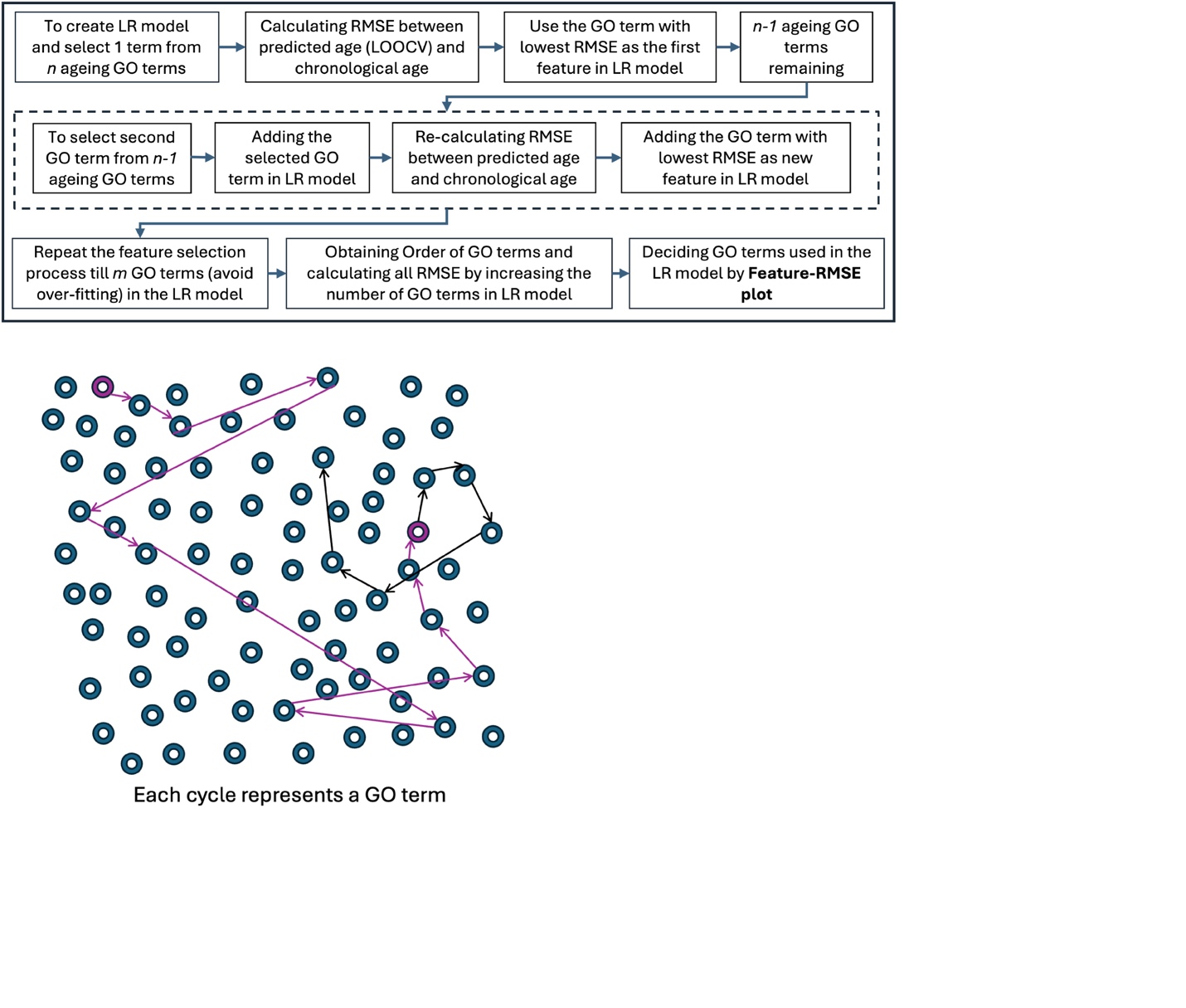

**Figure S2 The flowchart of Time Traversal Algorithm.** The flow of the algorithm (top) and the selection of the features (bottom).

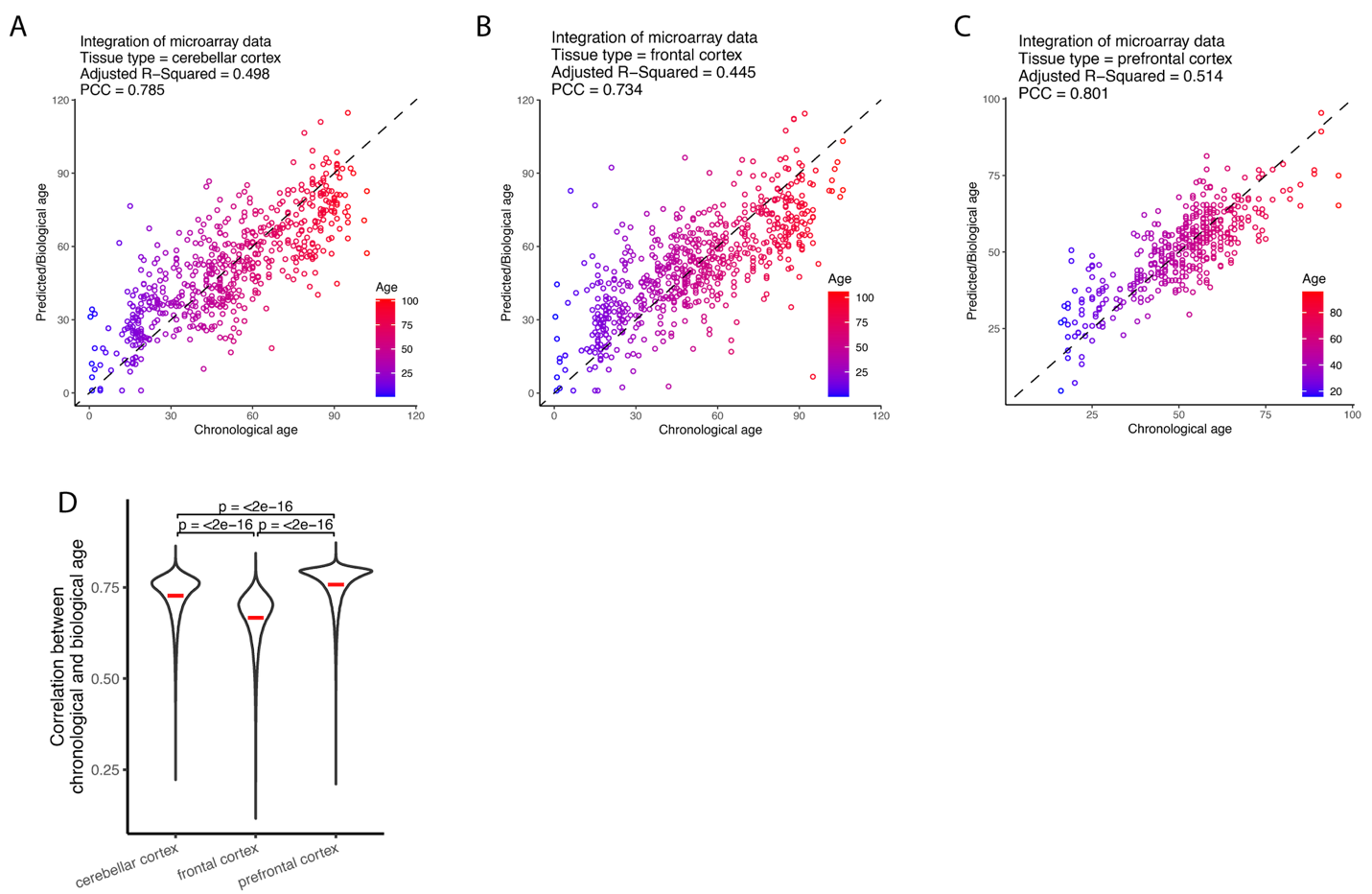

**Figure S3 The prediction of biological age in independent microarray data sets**

A-C) The LR models to predict biological age within one cell type in the integrated five microarray data sets.

D) By down-sampling of the samples within each sample type, the violin plot shows the ‘Pearson’ correlation coefficients.

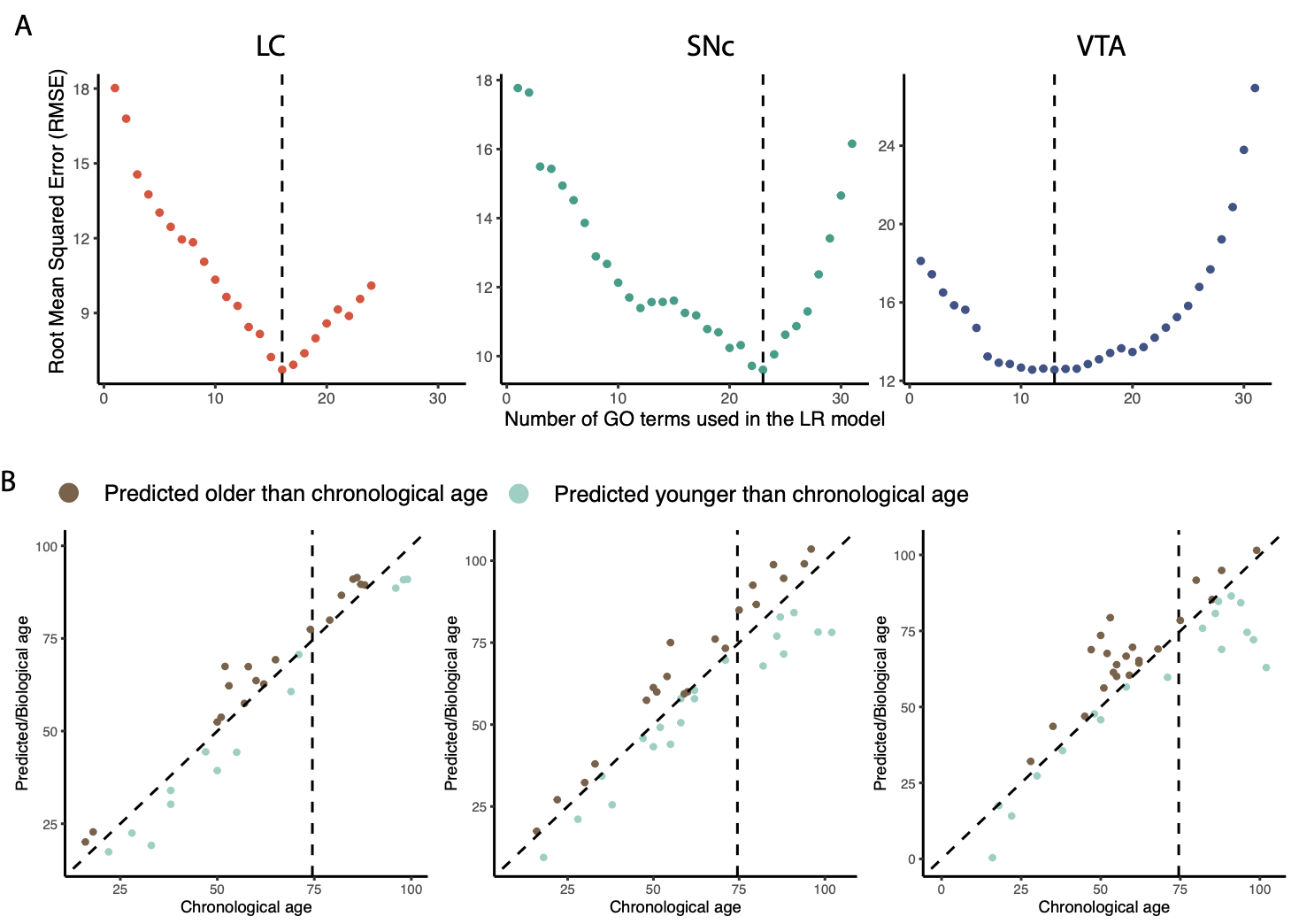

**Figure S4 The prediction of biological age within each sample type**

A) The feature-RMSE plots of the LR models in LC (left), SNc (middle) and VTA (right).

B) The samples were clustered into predicted younger and predicted older, than their chronological age, with the age of 75 years, indicated with a dotted line, shown for LC (left), SNc (middle) and VTA (right).

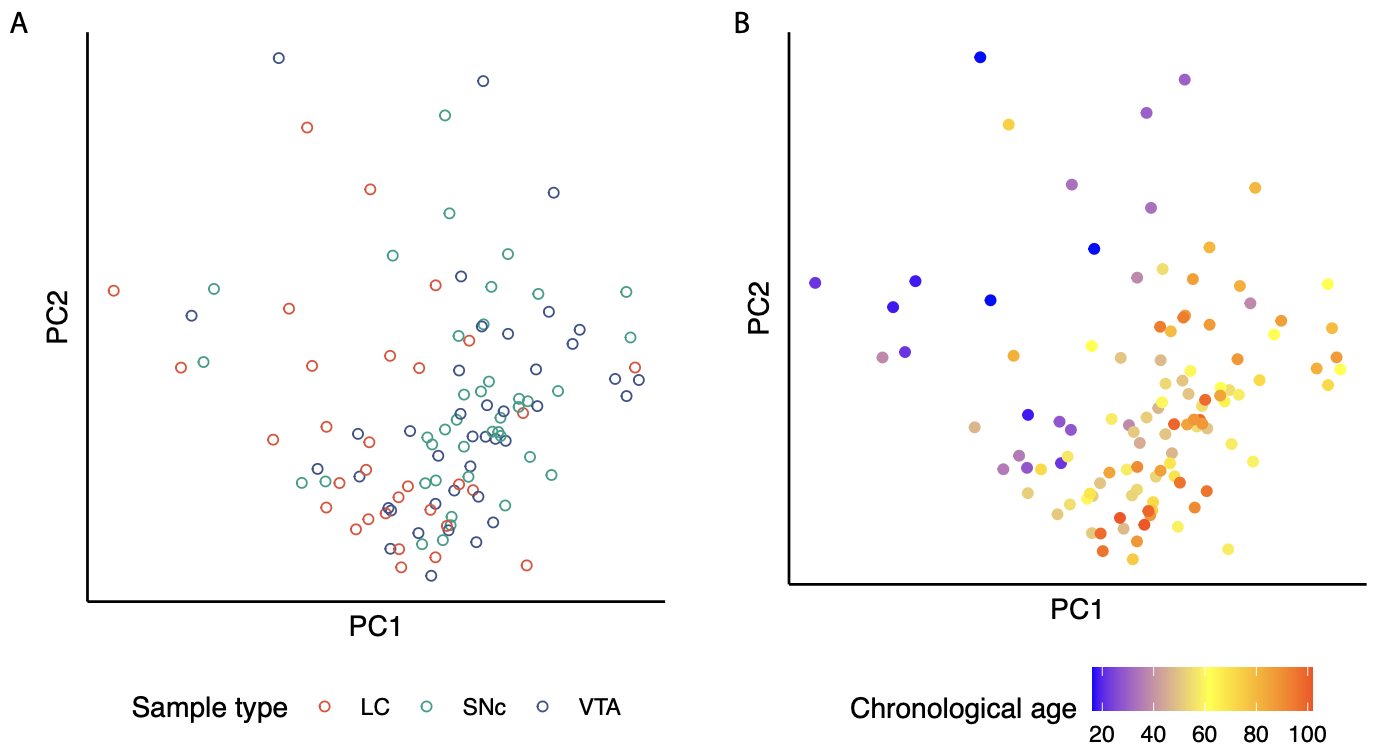

**Figure S5 The dimension reduction of the 111 samples by using 2764 genes.**

The PCs were calculated by using 2,764 ageing genes and the samples were colored by sample types (A) or by chronological age (B).

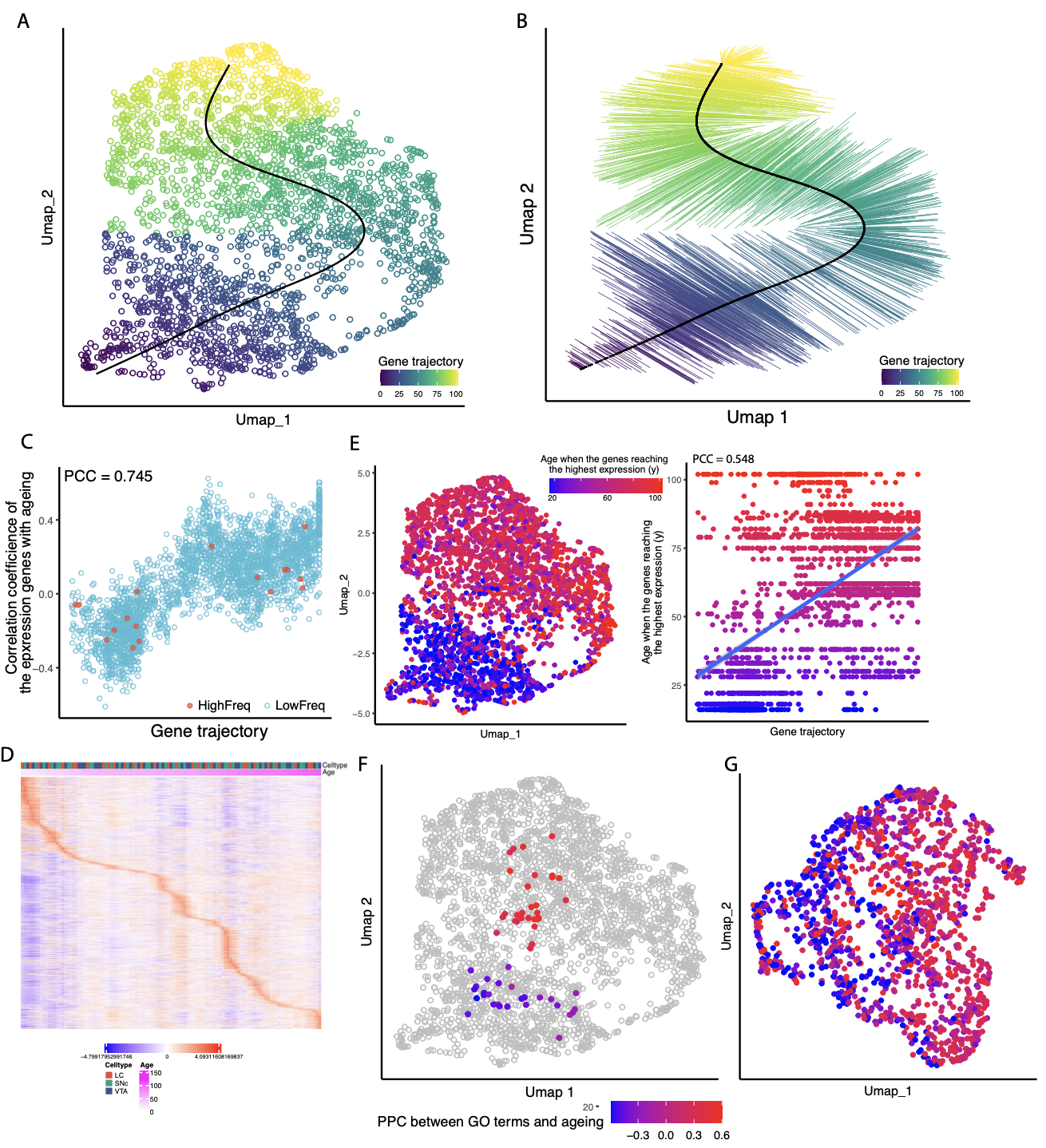

**Figure S6 The gene trajectory of the 2764 genes**

A-B) The 2,764 genes were projected into the UMAP by using the 111 samples collected by LCM-seq. The gene trajectory (A) of the genes was calculated based on the principal curve (B). C) The scatter plot shows the relation between gene trajectory and PCC with ageing.

D) The heat map shows the gene expression signature (details in Method) of the 2,764 genes.

E) The distribution of the values of gene expression signature in Umap (left) and gene trajectory (right). F) The colored dots show the average coordinates of the genes from the 59 GO terms. Gray dots are the 2,764 genes. Blue/Red dots represent the GO terms. G) The PCC values of the genes from the independent microarray data sets.

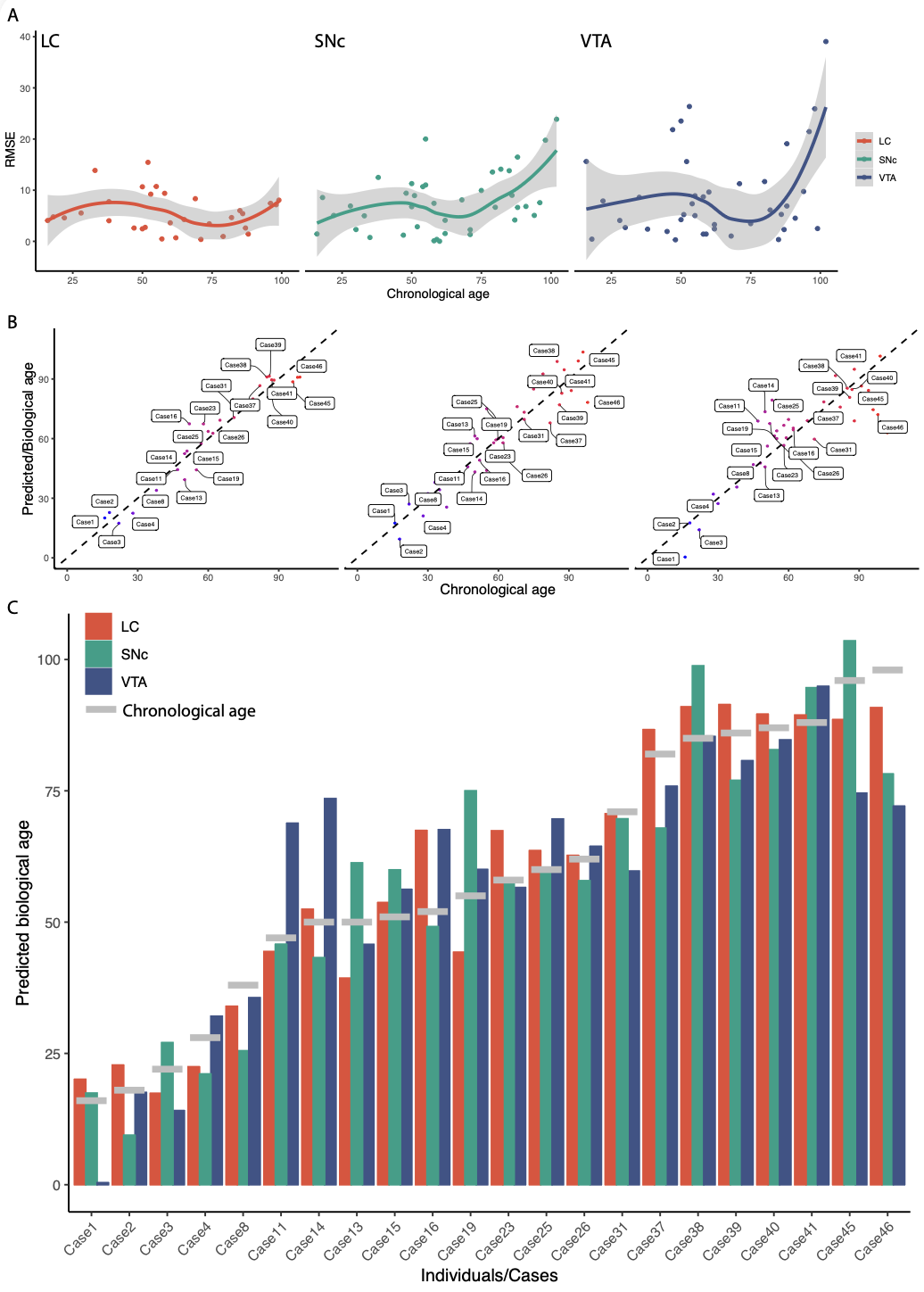

**Figure S7 The differences during the biological age prediction among LC, SNc and VTA.**

A) The RMSE with ageing in LC (left), SNc (mid) and VTA (right). B) The scatter plots show the samples of LC (left), SNc (mid) and VTA (right) originating from the same individuals (Cases).

C) The bar plot shows the predicted biological age of the sample of LC, SNc and VTA originating from the same individuals (Cases).

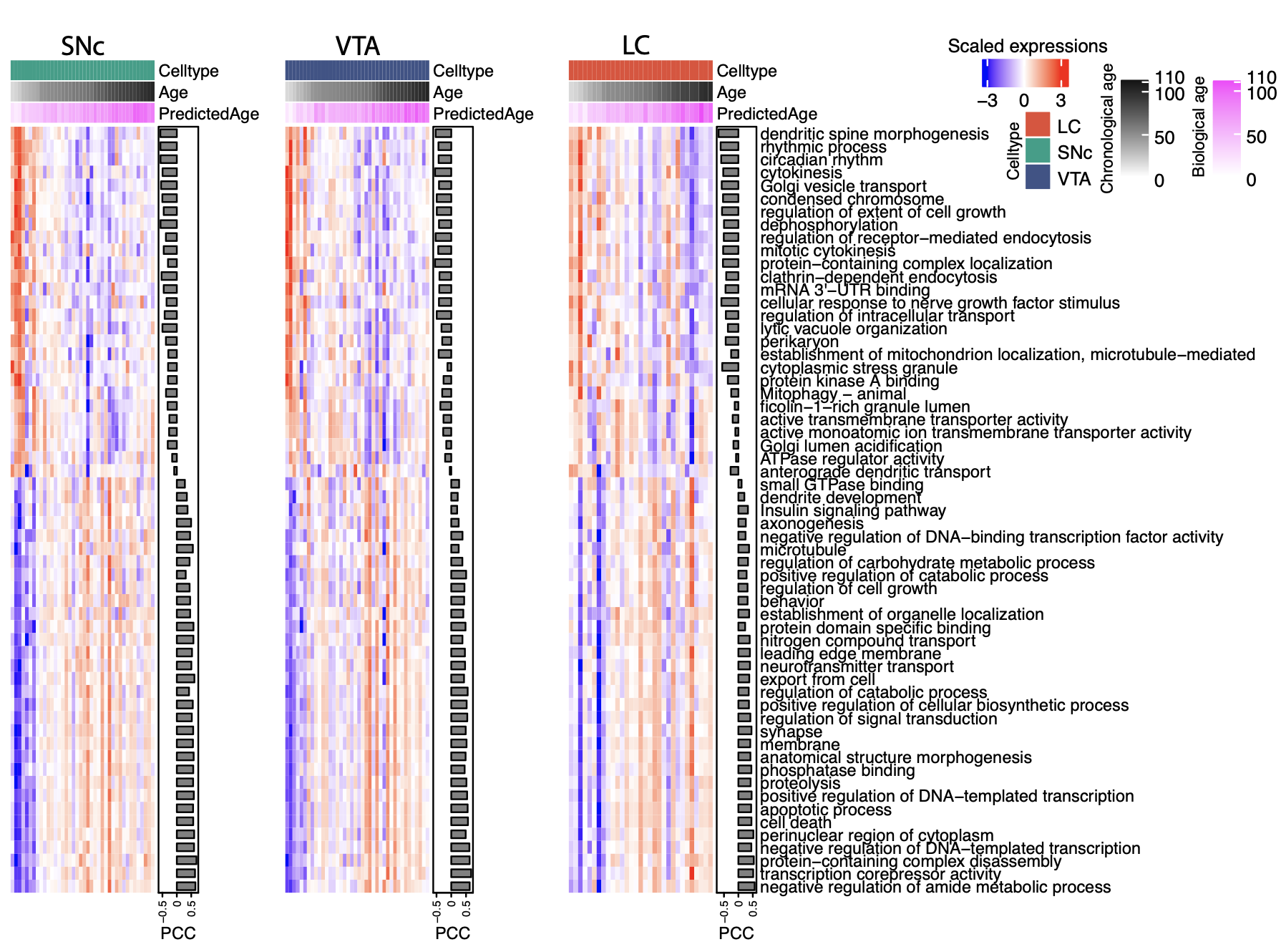

**Figure S8 The expressions of the ageing biomarkers**

The mean expressions of the genes from the 59 feature GO terms. The samples were ordered by the biological age from left to right. The side-by-side barplot is the ‘Pearson’ correlation coefficients between the mean expressions and the chronological age.

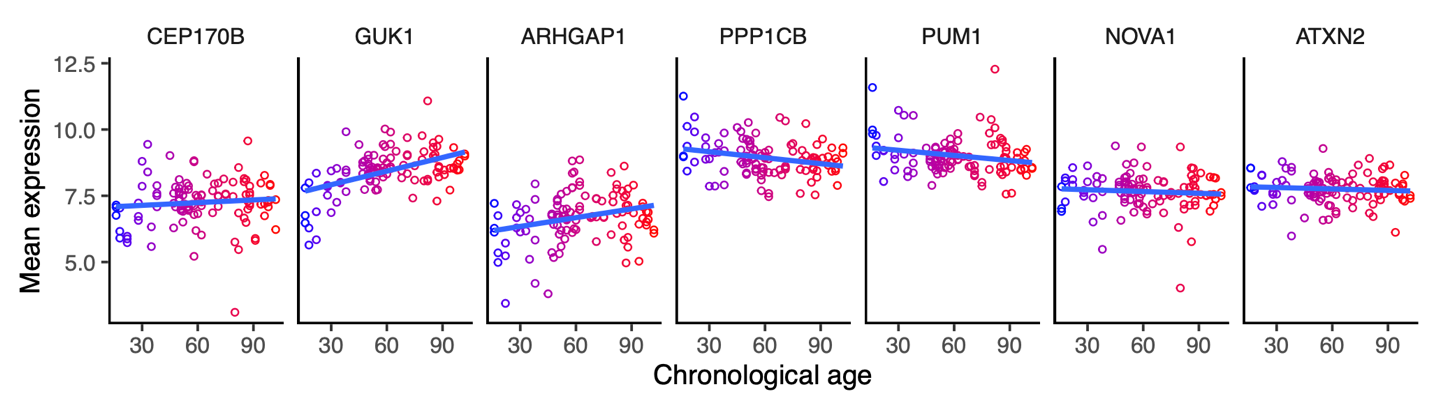

**Figure S9 The expression patterns of a set of ageing-regulated genes.**

The expressions of selected genes with regulation with age were normalized by DESeq2.

**Table S6 The ageing predictions in previous studies**

| **Published studies** | **Prediction** | **Features** | **PCC** | **R-squared** |
| --- | --- | --- | --- | --- |
| (Varshavsky et al., 2023) | GP-age | CpG sites with DNA methylation |  | Unadjusted R^2^ = 0.922 |
| (Horvath, 2013) | DNA methylation age | CpG sites with DNA methylation | 0.96 |  |
| (Hannum et al., 2013) | DNA methylation age | Methylation markers | 0.963 and 0.905 |  |
| (Soreq et al., 2017) | Biological age of the tissues in human brain | Cell-type specific genes |  | Unadjusted R^2^ = 0.35 ~ 0.58 |
| (Belsky et al., 2015) | Biological age | Biomarkers from NHANES | 0.38 with pace of ageing |  |
| (Husted et al., 2022) | Biological age of woman and man | Physical activity scales and quality of life | 0.86 and 0.81 | Unadjusted R^2^ = 0.73 and 0.65 |
| (Qiu et al., 2023) | Rescaling biological age (ENABL age) | Biomarkers from NHANES | 0.7867 and 0.7126 |  |
